# LRcell: detecting the source of differential expression at the sub-cell type level from bulk RNA-seq data

**DOI:** 10.1101/2021.08.10.455821

**Authors:** Wenjing Ma, Sumeet Sharma, Peng Jin, Shannon L. Gourley, Zhaohui Qin

**Affiliations:** Department of Computer Science, Emory University, 400 Dowman Drive, Atlanta, GA 30322, USA; Graduate Program in Neuroscience, Emory University, 1462 Clifton Road NE, Atlanta, GA 30322, USA; Department of Human Genetics, Emory University, 1365 Clifton Road, Atlanta, GA 30322, USA; Department of Pediatrics, School of Medicine, Emory University, 100 Woodruff Circle, Atlanta, GA 30322, USA; Yerkes National Primate Research Center, Atlanta, GA 30322, USA; Department of Biostatistics and Bioinformatics, Rollins School of Public Health, Emory University, 1518 Clifton Road NE, Atlanta, GA 30322, USA

**Keywords:** Cell-type enrichment, Cell marker genes, Differential gene expression

## Abstract

Given most tissues are consist of abundant and diverse sub cell-types, an important yet unaddressed problem in bulk RNA-seq analysis is to identify at which sub cell-type(s) the differential expression occur. Single-cell RNA-sequencing (scRNA-seq) technologies can answer the question, but they are often labor-intensive and cost-prohibitive. Here, we present *LRcell*, a computational method aiming to identify specific sub-cell type(s) that drives the changes observed in a bulk RNA-seq experiment. In addition, *LRcell* provides pre-embedded marker genes computed from putative single-cell RNA-seq experiments as options to execute the analyses. Using three different real datasets, we show that *LRcell* successfully identifies known cell types involved in psychiatric disorders and *LRcell* is more sensitive than even the leading deconvolution methods.

## Background

Finding genes that are differentially expressed (DE) between experimental conditions is a powerful approach to understand the molecular basis of phenotypic variation. However, most tissues are consisted of tens or even hundreds of diverse sub-cell types and DE may only occur in a small subset of these sub-cell types, which are relevant to the experimental condition. Bulk RNA-seq data alone is unable to reveal the sub-cell types that drives the DE.

The rapid development and proliferation of single cell technologies resulted in massive accumulation of single cell transcriptomics data (scRNA-seq) from diverse tissue types. These data reveal substantial variations in transcriptional regulation among different cell types and offer an unprecedented close-up view of the modifications underlying important biological processes including which cell types drive DE [1]. However, steep cost and complicated protocols prevent the wide-spread adoption of scRNA-seq.

In this study, we propose a novel computational tool named *LRcell*. Given the result from a bulk RNA-seq DE study, the goal of *LRcell* is to delineate which sub-cell type(s) of the tissue underwent substantial changes between the two experimental conditions. Exploiting cell type-specific marker genes identified from generic scRNA-seq available from publicly available data repositories, *LRcell* achieves the goal by surveying the enrichment of marker genes across all sub-cell types in the tissue (Figure 1). Thus, no scRNA-seq experiment matching the bulk RNA-seq experimental condition is needed. When applying *LRcell* to a diverse panel of bulk RNA-seq DE experiments, we successfully identify known cell types involved in the pathogenesis of psychiatric disorders as well as produce testable new hypotheses that have the potential to produce fresh new biological insights.

**Figure 1.**
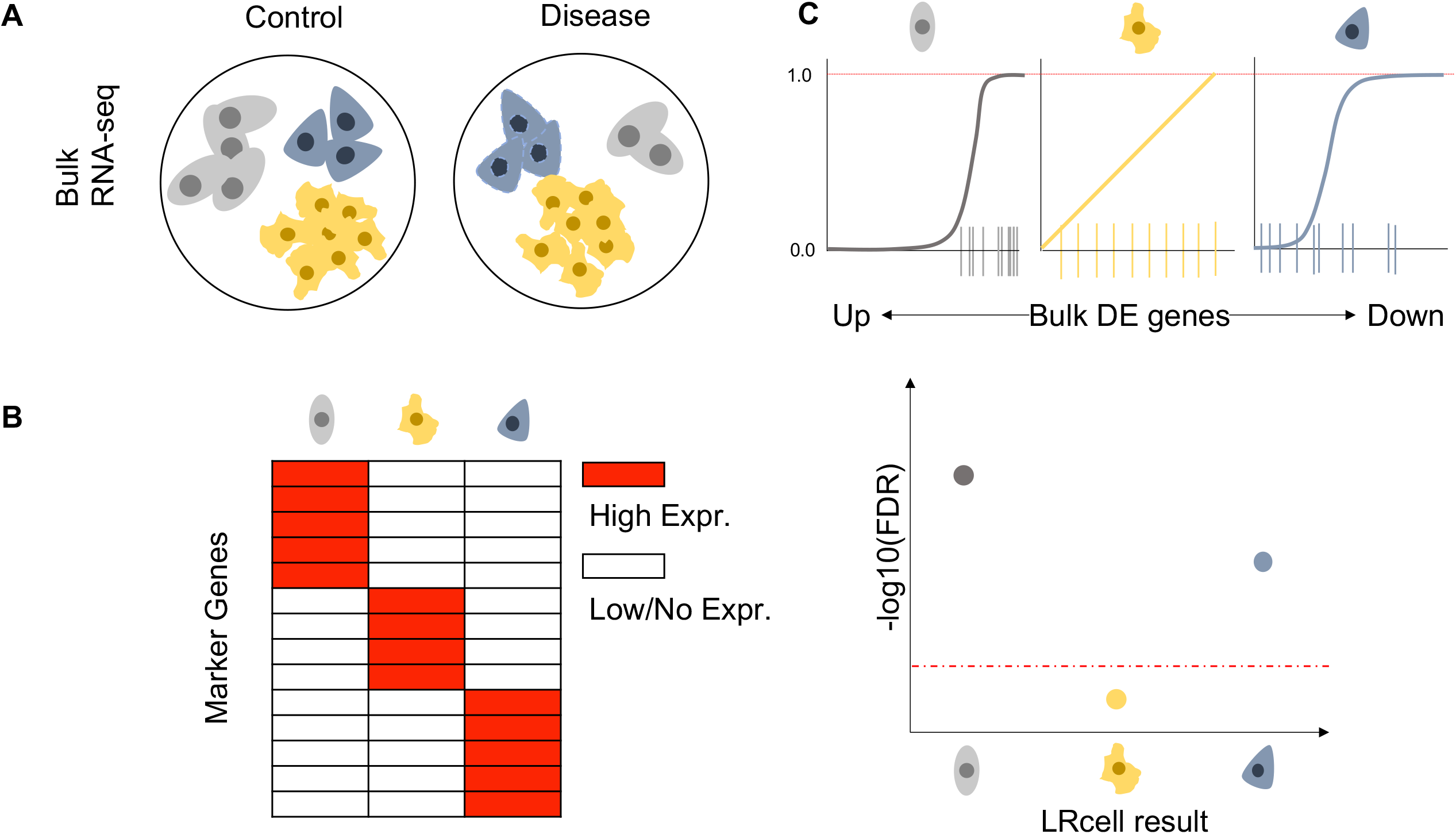
Overview of *LRcell* workflow. (A) Bulk RNA-seq experiment design with control and disease conditions. (B) Illustration of a marker gene list derived from scRNA-seq experiment. (C) (upper panel) Illustration of how the distribution of marker genes on bulk DE genes may affect the regression; (lower panel) Illustration of the FDR rate for each cell type in which the grey cells and dark blue cells are most likely involved in the condition change.

## Results and Discussion

In this work, we collect and curate a compendium of marker genes from multiple published scRNA-seq datasets. We then conduct *LRcell* analysis on multiple bulk RNA-seq DE experiments to demonstrate its utility.

### 3.1 Marker gene collection and sources

Similar to a collection of gene set for Gene Set Enrichment Analysis (GSEA) [2], *LRcell* requires a compendium of high-quality cell type marker genes. Currently, *LRcell* package provides users with multiple pre-loaded marker gene sets from human blood, human brain and mouse brain (Figure 2.A), computed from scRNA-seq datasets. Additionally, *LRcell* package offers external cell markers collected by Molecular Signatures Database (MSigDB) [3]. The external makers all originate from human species including midbrain, cord blood, ovary and skeletal muscle. We store all cell-type specific marker gene sets into another R Bioconductor ExperimentHub package named *LRcellTypeMarkers*. Additional marker gene sets are being tested and will be added to the collection.

**Figure 2.**
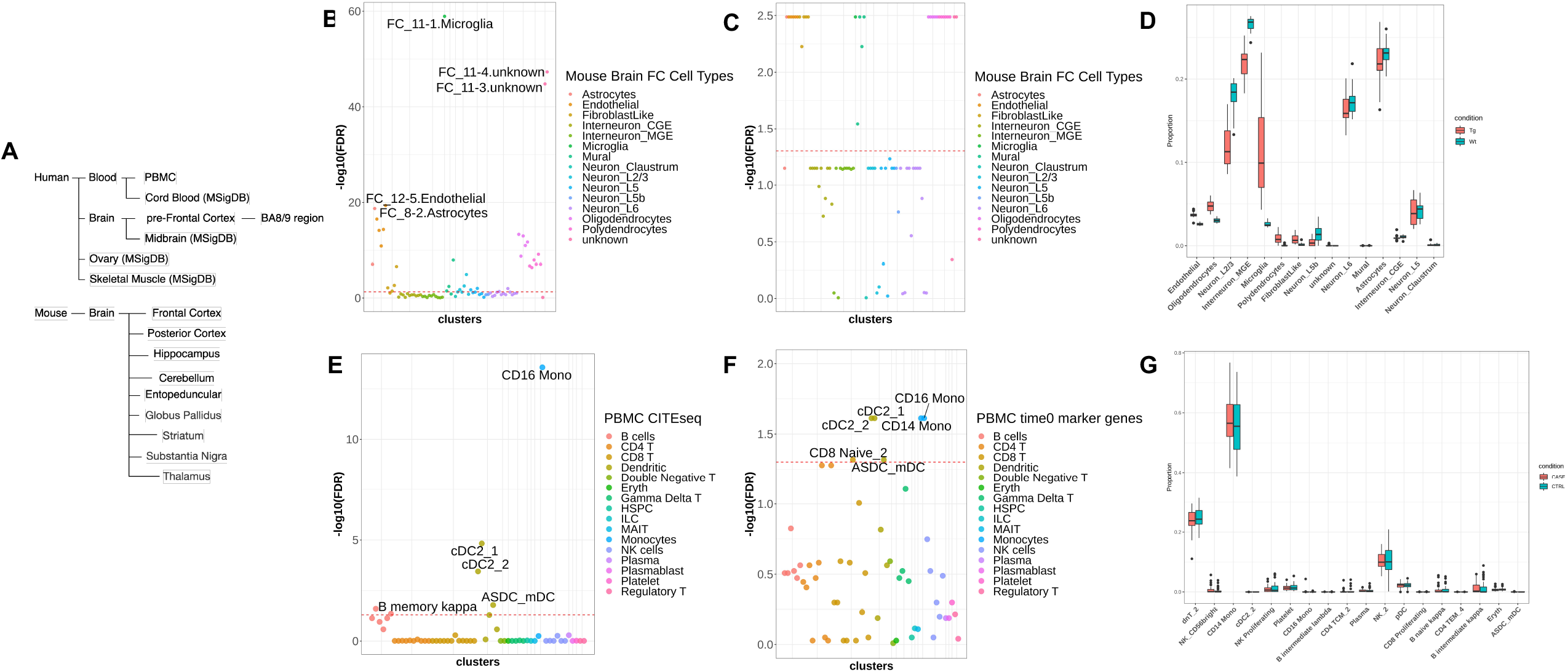
*LRcell* datasets and *applying LRcell* to real cases. (A) summary of the all tissue types in which marker genes have been pre-embedded in *LRcell*. (B) *LRcell* result of mapping the bulk neurodegenerative dementia DE genes to the mouse brain frontal cortext (FC) region . Microglia is shown as the most significant cell type. (C) GSEA result of mapping the bulk neurodegenerative dementia DE genes using the same marker genes for *LRcell* (mouse brain FC) as input. We see numerous dots have the same significance level which is difficult to interpret. (D) Cell type proportions for control and disease samples calculated by MuSiC. Each box contains 17 individuals. The x-axis is ordered by the t-test significance between the two conditions. (E) *LRcell* result of mapping bulk PTSD DE genes to human PBMC. CD16+ Monocytes is shown as the most significant cell type. (F) GSEA result of mapping bulk PTSD DE genes to human PBMCusing the same marker genes for *LRcell* (human PBMC) as input. Again, we see numerous dots have the same significances. (G) Cell type proportions for control and disease samples calculated by MuSiC. The x-axis is ordered by the t-test significance between two conditions. As shown in the figures, MuSiC failed to identify all cell type proportions as there are plenty of 0s starting from ASCD_pDC cell type and the CD14+ Monocytes are wrongly inferred.

### 3.2 Microglia highly enriched in Neurodegenerative Dementia

In a recent neurodegenerative dementia study, Swarup and colleagues contrasted TPR50 mice expressing tau mutant with wild type mice using bulk RNA-seq in order to identify gene networks mediating dementia [4]. To identify the cell type(s) most involved in the condition, we apply *LRcell* to the DE gene list using pre-embedded marker genes from adult mouse Frontal Cortex (FC) region [5]. From *LRcell* result, we observed that Microglia show up as highly significant (Figure 2.B) which is concordant with previous studies [6]. Additionally, the FC_11-3.unknown and FC_11-4.unknown sub cell types also show high level of significance. No annotation is available for these two cell clusters in the original publication. However, pair-wise comparison of marker genes among all cell clusters reveal that these two unknown cell clusters have considerable overlaps with the FC_11-1, which is also a Microglia cell type (Figure S1. A), which explains the pattern we observe.

### 3.3 CD16+ Monocytes highly enriched in Post-Traumatic Stress Disorder (PTSD)

In a recent study, Breen and colleagues conducted a bulk whole-transcriptome study using peripheral blood leukocytes collected from U.S. Marines, among which some developed PTSD post-deployment [7]. Using this dataset, we generate a list of DE genes that show significant difference between the PTSD group and the control group at the pre-deployment timepoint.

Using human marker genes derived from a single cell transcriptomic study on PBMC [8], *LRcell* analysis finds that cells annotated as CD16+ non-classical Monocytes shows up as the most significant among all cell types in PBMC (Figure 2. E). Our finding makes biological sense because as stated in previous studies [9], heterogeneity exists in monocytes distinguished by CD16 surface proteins and non-classical monocytes have been validated to regulate immune responses in trauma [10,11].

### 3.4 Marker genes from different region or timepoints

To apply *LRcell*, an important question is that which marker gene sets to use, i.e., how to select single cell RNA-seq data where the source of the tissue match the tissue type profiled in the bulk transcriptomic study. This is particularly important for complex tissues such as brain. To address this issue, we use the neurodegenerative dementia study [4] as an example, which contains information from four brain regions: cortex, hippocampus (HC), cerebellum (CB) and brain stem. Brain stem is excluded from our analysis due to the lack of marker gene information from that region of the brain.

To understand how marker genes vary across brain regions, we first define marker genes in all regions of the brain to explore their spatial pattern (Figure S1. B; Figure S1. C). We observe that glia cells, such as Astrocytes, from different regions have higher number of overlapping marker genes which indicates the homogeneity of glia cells across the brain. In contrast, neurons and interneurons share very few marker genes across different brain regions. We then apply pre-embedded adult mouse brain marker genes from FC, HC and CB to bulk DE genes obtained from cortex, HC and CB respectively (Figure S3). We observe that Microglia cells are highly enriched in all three brain regions whereas Astrocytes are particularly highly enriched in CB (Figure S3). Especially when applying CB marker genes to CB bulk DE experiment, we notice that one sub-cell type of Astrocytes is highly enriched compared to others. Our observations demonstrate that selected cell types are heterogeneous spatially, meaning marker genes are highly specific not only for the cell type, but also which region the cell belongs to. Because of this finding, it is highly desirable to run *LRcell* using marker genes of cell types located in closely matched brain regions.

We are also curious whether marker genes curated from scRNA-seq experiments conducted on non-normal samples is acceptable as the reference. To address this question, we use data from the HIV vaccine study [8], we observe that the expression of cell-type specific marker genes is mostly consistent across different timepoints within the same cell type (such as CD8 cells), and distinct across different cell types (Figure S2.A; Figure S2.B). For example, B sub-cell types share a considerable number of marker genes across timepoints, while sharing fewer with other cell types. Thus, although default marker genes used in *LRcell* are collected from control samples, we believe that marker genes identified from non-normal samples are acceptable when scRNA-seq data from normal samples are not available.

### 3.5 Comparison to GSEA

GSEA is a powerful tool to determine whether a pre-defined gene set show concordant shift in expression when comparing two biological conditions. One could potentially replace *LRcell* with GSEA to identify DE-driving cell types, by treating cell-type specific marker genes as pre-defined gene sets. To compare performance of the two methods, we repeat the analyses done in section 3.1 and 3.2 using GSEA. The GSEA result from the neurodegenerative dementia mouse model (Figure 2C) yields several equally significant cell types including Astrocyte, Endothelial, Microglia, Mural, Oligodendrocyte and Polydendrocyte. The tied significances lead to difficulties in determining which cell type(s) potentially participated in dementia pathogenesis. Similar pattern is observed in the GSEA result on PTSD study (Figure 2F) which shows that Monocytes, Dendritic cells and some T sub-cell types are equally enriched. Based on the above observations, we conclude that *LRcell* is more effective than GSEA to identify sub cell types that are most impacted by the condition change in bulk DE experiments.

### 3.5 Comparison to MuSiC

A plethora of deconvolution methods has been developed to infer the proportions of different sub cell types from bulk transcriptomic data [1][12][13][14][15]. By deconvoluting individual bulk samples and comparing their proportions, one can identify both cell-type proportion and gene expression changes. Hence it is of interest to compare *LRcell* with such a bulk deconvolution strategy. For comparison, we apply MuSiC [13], a deconvolution method that have shown favorable performance, to the mouse neurodegenerative dementia study [4] on both conditions. Because some layers of neurons are predicted to have almost zero proportion (Figure 2D) when using all 81 sub-cell types, we merge the original sub-clusters into 15 major cell types in order to achieve a better representation. Despite this, MuSiC does not detect significant difference in Microglia or Astrocyte in terms of their proportions between the two conditions. When applied to the PTSD study, using the original cell cluster annotations, MuSiC shows that most of the T sub-cell types have zero proportion and the proportion of CD14+ Monocytes is up to 60% (Figure S2G). In contrast, *LRcell* produces more sensible results because it is not limited by the number of cell types as it can detect the subtle differences among sub-cell types.

### 3.5 Discussions

Detecting transcriptional activity changes at the individual cell type level, especially their modifications in disease samples, is crucial for understanding the mechanisms of diseases development. Deconvolution-based computational tools have been developed to dissect cell type proportions from bulk gene expression profiles. The computationally inferred proportions can then be used for comparison. However, when apply to real data, such a strategy often encounters numerical stability problem and unable to handle large number of sub-cell types. In this study, we propose a novel alternative strategy named *LRcell, LRcell* conducts enrichment analysis of cell-type specific marker genes among the top (or bottom) DE genes identified by bulk transcriptome studies. Cell types that show the most enrichment is likely to play an important role in the condition alteration. When applying to real datasets, we found *LRcell* can successfully identify the involvement of the Microglia and Astrocytes in neurodegenerative dementia and rare Monocytes in PTSD.

In spirit, *LRcell* operates similarly as GSEA. But *LRcell* is much more sensitive to minor differences in marker genes of sub-cell types, which indicates its potential to detect changes in sub-cell types caused by disease conditions. Additionally, when compared to existing bulk deconvolution methods, *LRcell* is more stable in its ability to handle the similarities among sub-cell types. Thus, *LRcell* enables researchers to glean new biological insights from the bulk transcriptomics experiments with no need of redo the experiment using single cell technology. We are currently applying *LRcell* to a diverse set of clinical studies (Sharma, personal communication) to generating more biological insights.

To enable straightforward comparison, currently, we select a fix number of 100 marker genes from each sub-cell type. Understandably, the number of marker genes for different cell types varies, it is desirable to allow flexibility in choosing the number of marker genes based on the transcriptomic patterns across cell types. However, different numbers of marker genes post challenge for conducting enrichment analyses fairly across all cell types. This will be investigated in our future studies.

*LRcell* currently provides embedded marker genes from human blood, human brain and mouse brain calculated from scRNA-seq experiments along with markers from 66 cell types in four tissues (midbrain, cord blood, ovary, and skeletal muscle) adopted from MSigDB. We are working to include more tissue types in the future releases of *LRcell* which will make it more widely applicable.

## Conclusions

In summary, we develop *LRcell*, an R Bioconductor package for identifying sub-cell type(s) that drive the changes observed in bulk comparative transcriptomic studies, taking advantages of newly emerged scRNA-seq data. The rationale of *LRcell* is that we believe marker genes of the modifying cell types tend to be enriched toward the top (or bottom) of the DE gene lists. We conduct comprehensive surveys applying *LRcell* across various experimental conditions, and successfully identify cell types that play important roles in neurodegenerative dementia and PTSD. Hence, we believe *LRcell* can provide researchers important and new biological insights in terms of the source of the biological changes at the sub-cell type level, without the need of conducting costly and laborious scRNA-seq experiments.

## Methods

### Basic Assumptions

The goal of *LRcell* is to identify the most affected sub-cell type(s) during the transition of experimental conditions using only bulk transcriptomic data. Based on the assumptions that cell-type specific marker genes of key cell types tend to be over-represented among the significant DE genes in bulk transcriptomic studies, *LRcell* can discover which cell type(s) is involved in certain disease or condition change. In recent years, computational methods have been developed to deconvolve bulk RNA-seq data to delineate cell-type proportion changes, which could be borrowed to answer the same question. However, whenever there are ten or more sub-cell types, the results from deconvolution methods become unreliable. In contrast, *LRcell* enables comparison across many more cell types which is important for complex tissues such as brain.

The development of *LRcell* is inspired by LRpath [16], which is designed for linking experimental changes to biological pathways or a pre-defined gene set. In *LRcell*, we treat cell-type specific marker genes as gene sets and calculate the enrichment of each cell type when comparing two biological conditions. We believe that the most enriched cell type(s) is highly likely to play an important role in the experimental condition change.

### scRNA-seq Data Preprocessing

In this study, we include marker genes from mouse whole brain, human prefrontal cortex (pFC) and human PBMC, along with 66 cell types’ markers from four tissues (midbrain, cord blood, ovary and skeletal muscle) adopted from MSigDB. For each scRNA-seq dataset, we first retrieved raw read count matrix. Next, we filtered out low-quality cells and genes and apply column-wise normalization and log transformation on the data.

The mouse whole brain scRNA-seq dataset [5] produced using the Drop-seq technology [17], contains nine brain regions from adult mice. The data provided has already been pre-filtered by the authors. For cell types other than neurons, we directly utilized the information provided on the study website. For neurons and interneurons, we curated the sub-cell types following the original study.

The human pFC scRNA-seq dataset [18], produced by 10X Genomics Chromium, is derived from the pFC region (specifically BA9). The dataset contains two conditions: healthy controls and major depressive disorder (MDD). We split the data matrix into two parts and filtered out cells expressing less than 10 genes and genes expressed in less than 10 cells respectively. We also filtered out mitochondrial, ribosomal genes, and genes from annotation clusters (Astros_1, Mix_1, Mix_2, Mix_3, Mix_4, Mix_5, Inhib_4_SST).

The PBMC dataset [8], generated by CITE-seq technology [19], is derived from an HIV vaccine trial study which involves eight volunteers at three timepoints: immediately before, three days and seven days after the vaccine. The study contains 161,764 cells in total. To accelerate the marker gene selection, we separated the count matrix according to the time label and filtered out low-quality cells and genes (mitochondrial, ribosomal genes and those expressed in less than 1,000 cells). The cluster annotated as “Doublet” was filtered out.

### Marker Gene Selection

After obtaining the log-normalized gene expression matrix along with high-quality sub-cell type clusters, we calculated the enrichment scores for each cluster (sub-cell type) using the marker gene selection method described in Marques et al[20]. The cluster-specific gene enrichment is defined as the average gene expression levels of cells in that cluster divided by the average gene expression levels in all cells. The enrichment score is adjusted by introducing a penalty representing the fraction of cells in that cluster expressing the marker gene. Combined, this score allows the identification of genes with cluster-specific high expression values to be selected as marker genes. The description below is adapted from the original publication.

Suppose there are a total of *M* genes, *L* different clusters each with *N*_*j*_ cells, and the total number of cells is *N*. Let *E* = {*E*_*ijk*_} represent the gene by cell read count matrix. Here *j*=1 *i* = 1, …, *M, j* = 1, …, *L, k* = 1, …, *N*_*j*_ and 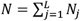. The overall average expression of the ith gene across all cells is defined as:

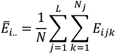

The average expression of gene *i* in the *j*th cluster is defined as:

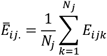

The enrichment for gene *i* in the *j*th cluster as:

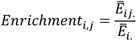

Next, we consider the proportion expressing the gene *i* in the *j*th cluster as:

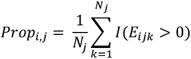

The *I*(·) is an indicator function.

The enrichment score for gene *i* in the *j*th cluster is computed as:

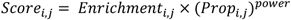

where “power” is a hyperparameter to be tuned manually to control the penalization for the cell cluster proportion term. The power parameter is set to 1 throughout this study. After calculating the weighed gene enrichment scores in each cluster, we ranked genes based on the scores and selected the top 100 genes as the marker genes for each cluster.

### MSigDB Marker Genes

We downloaded cell marker gene sets from MSigDB category C8 – cell type signature gene sets. Since not all tissue types are suitable for *LRcell*, we applied the following criteria to select tissues: 1) non-fetal tissues; 2) have more than 8 (sub-) cell types; 3) minimum number of marker gene greater than 50; 4) median number of marker genes greater than 80. In the end, four tissue types, the midbrain, cord blood, ovary and skeletal muscle remain.

### Bulk RNA-seq Data Preprocessing

The raw count of mouse bulk RNA-seq study on neurodegenerative dementia was downloaded from GEO (Accession number: GSE90693). DE analysis is performed using DESeq2 [21] to obtain DEGs in each brain region.

The raw count of bulk RNA-seq study on PTSD was downloaded from Recount2 [22]. We extracted out the experiment contrasting PTSD cases and healthy controls with timepoint of pre-employment and performed DESeq2 to obtain DEGs.

### LRcell Analysis

*LRcell* is inspired by LRpath, which was designed for identifying sets of predefined gene sets that show enrichment with differentially expressed transcripts in microarray experiments. *LRcell* uses logistic regression or linear regression to assess whether marker genes (as defined in the Marker Gene Selection sub-section above) of a specific cell type are more likely to be DE genes in a particular bulk RNA-seq study. The linear regression option is added to handle the continuous enrichment scores of marker genes. Users can choose accordingly. To facilitate our analysis, we assume that the major sub cell types of the tissue their marker genes are known *a priori*.

We apply *LRcell* to each cell type independently. The required input includes a list of DE genes ranked by the level of significance, and a set of marker genes for each cell type.

Then *LRcell* runs a logistic regression as

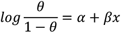

and

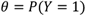

In which *Y* = 1 denotes that gene is a marker gene and *Y* = 0 otherwise. Hence *θ* represents the chance that the gene is a marker gene. We use −log (*p* − value) as the explanatory variable *x*. Whether a specific cell type is involved in the experimental condition change is evaluated by testing the null hypothesis that *β* = 0 against the alternative that *β* ≠ 0 using the Wald test. Typically, we run *LRcell* on all sub cell types found in the tissue to see which sub cell type(s) drives the changes.

Similar to logistic regression, linear regression directly performs

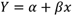

where *Y* indicates the enrichment scores of genes. Same as logistic regression, the p-value can be obtained from testing the null hypothesis that *β* = 0 against the alternative that *β* ≠ 0 using the t-test.

Once the p-values are obtained, we calculate False Discover Rate (FDR) using p.adjust() function in R to adjust p-values with Benjamini-Hochberg method.

### Input and Output

*LRcell* requires two inputs: (1) a ranked list of genes with DE p-values in a bulk RNA-seq experiment; (2) sets of marker genes from all sub-cell types of the bulk tissue acquired from scRNA-seq datasets *a priori* or from MSigDB C8—cell type signature gene sets. For those cell markers derived from scRNA-seq datasets, we offer choices for users to choose between species as human or mouse and the region indicates the specific brain region or PBMC. For MSigDB cell markers, we store the marker genes into the *LRcellTypeMarkers* packages which can be easily downloaded. When running LRcell() function, the Logistic Regression (LR) option is set as the default, while users can also set the method option as LiR if Linear Regression is desired. For linear regression, an enrichment score is needed as input for each gene whereas gene sets are sufficient for logistic regression. For MSigDB cell type signature gene set, LR option is recommended as there is no enrichment score information available. For customized input, i.e., a scRNA-seq data, we offer a LRcell_gene_enriched_scores function which takes the read counts matrix and cell annotation as input to generate enrichment scores for genes in each cell type. For further subsetting, get_markergenes can be used for generating marker genes for more specific sub cell types.

The output is a list of significance p-value (or FDR), one for each sub cell type. For visualization, *LRcell* produces Manhattan plot, which can be drawn through plot_manhattan_enrich function. We also provided a plot (plot_marker_dist function) indicating where certain cell-type specific marker genes locate on the bulk DE genes. The bulk DE genes are sorted using -log10(p-value) × sign(log2FoldChange) which could potentially give information on both up-/down-regulated directions. More detailed information about *LRcell* is available at http://bioconductor.org/packages/release/bioc/html/LRcell.html.

*LRcell* requires an R version beyond 4.1 and a prerequisite installation of BiocManager.

## Declarations

### Ethics approval and consent to participate

Not applicable.

### Consent for publication

Not applicable.

### Availability of data and materials

The datasets analyzed during the current study are available in the Gene Expression Omnibus (GEO) with following accession numbers: mouse whole brain [5] (GSE116470), human pFC [18] (GSE144136), human PBMC [8] (GSE164378), neurodegenerative dementia study [4] (GSE90696) and PTSD study [7] (GSE64814). The R package is freely available on Bioconductor (https://doi.org/doi:10.18129/B9.bioc.LRcell) and the external marker genes are stored in another R package named *LRcellTypeMarkers* on Bioconductor (https://doi.org/doi:10.18129/B9.bioc.LRcellTypeMarkers).

## Author’s contributions

ZQ and WM conceived the idea and supervised the study. WM realized the method and developed the R package. WM analyzed the real data and conducted the computational experiments. SS provided guidance on validating biological experiments and tested the package. WM and ZQ summarized the results and wrote the manuscript. All authors have read and approved the manuscript.

## Figures

**Figure S1:**
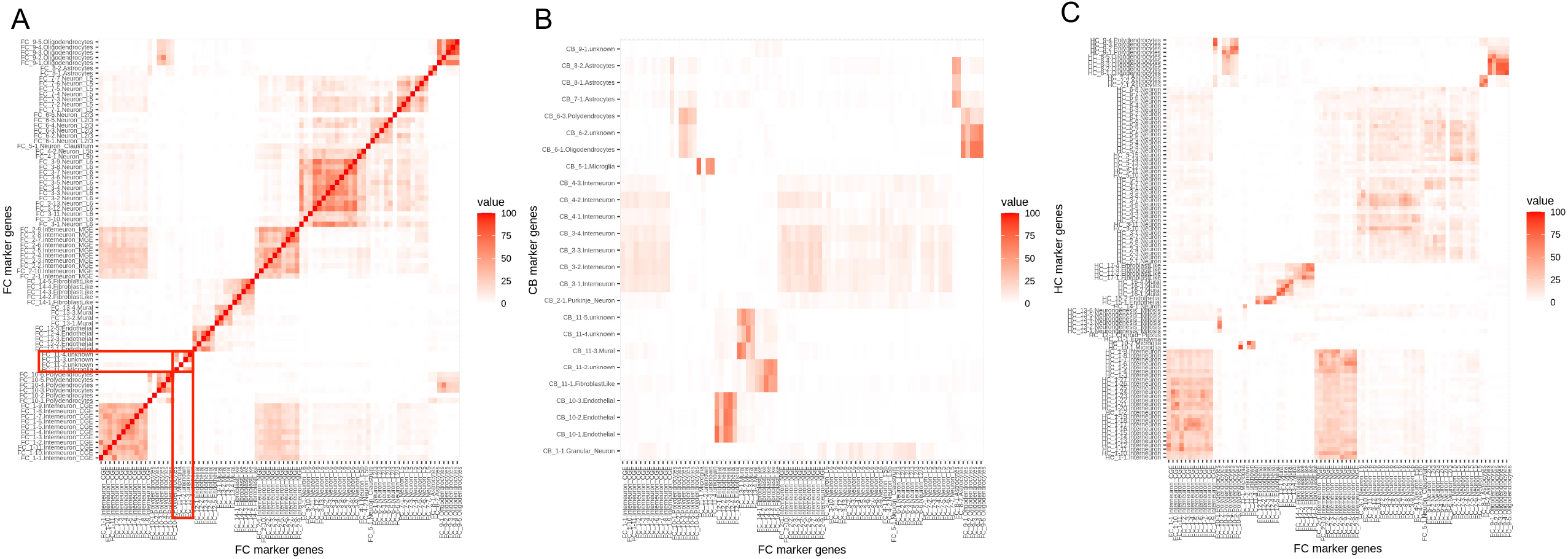
Marker genes overlap between different brain regions. Top 100 marker genes are selected for each cell type, thus, the maximum overlap in these figures is 100. (A) heatmap illustrating the overlap of marker genes among cell types within the Frontal Cortex (FC) region derived from mouse whole brain scRNA-seq dataset. The highlighted area describes the overlap between FC_11-3.unknown, FC_11-4.unknwon and FC_11-1.Microglia as an illustration for the similarity between these three sub-cell types. (B) Heatmap illustrating the overlap of marker genes among cell types within the FC and cell types within the Cerebellum (CB). (C) Heatmap illustrating the overlap of marker genes among cell types within the FC and cell types within the Hippocampus (HC). As shown in (B) and (C), the overlaps between cell types belong to different brain regions are lower than overlaps between cell types belong to the same brain region.

**Figure S2:**
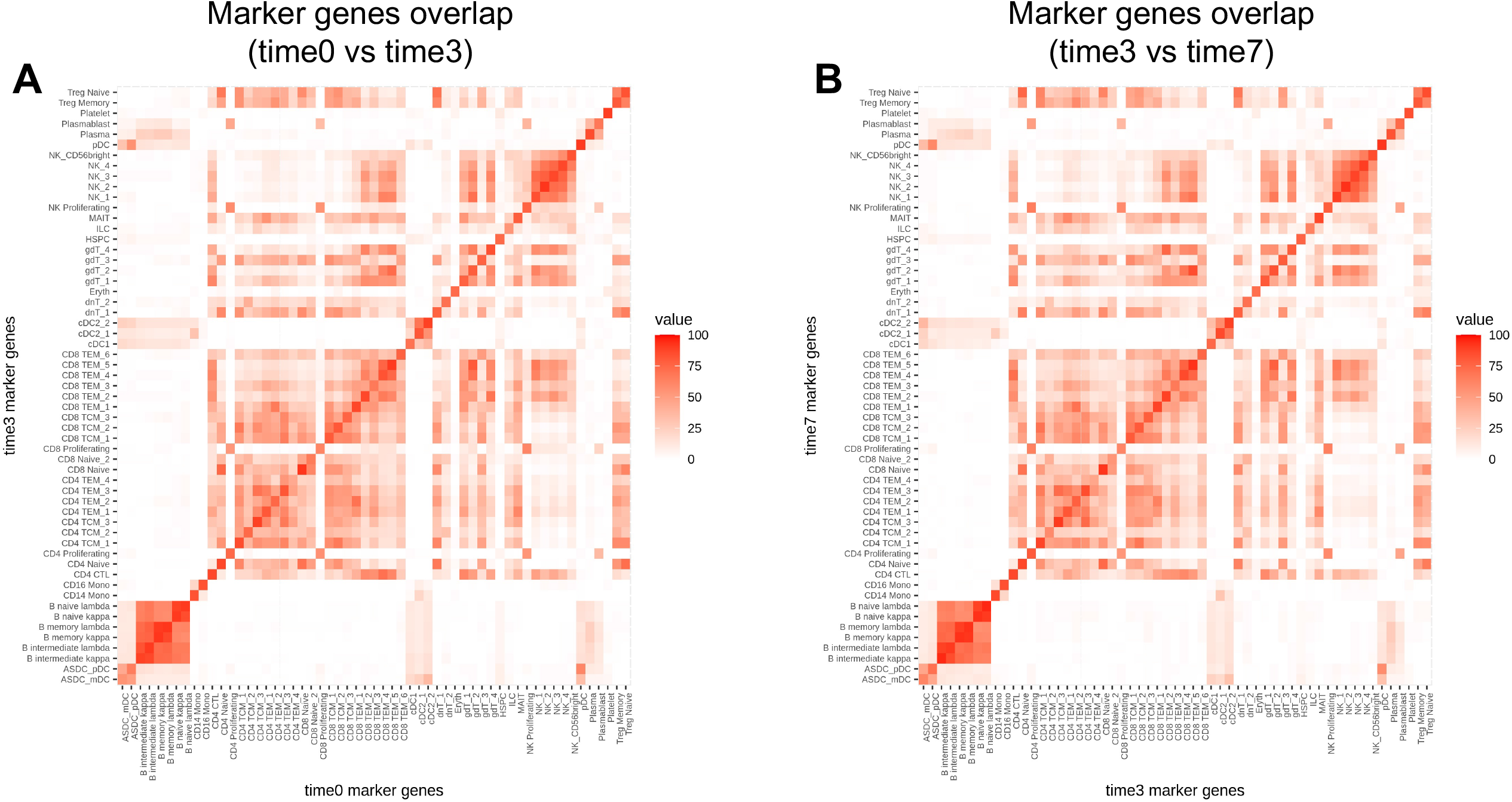
Marker genes overlap across time points from PBMC dataset. Top 100 marker genes are selected for each cell type and the maximum overlap in figures is 100. (A) Heatmap illustrating the overlap of the cell-type-specific marker genes between time point 0 and time point 3. (B) Heatmap illustrating the overlap of the cell-type-specific marker genes between time point 3 and time point 7. As shown in the figure, the diagonal pattern indicates the marker genes selected are highly cell-type-specific and shared across time points.

**Figure S3.**
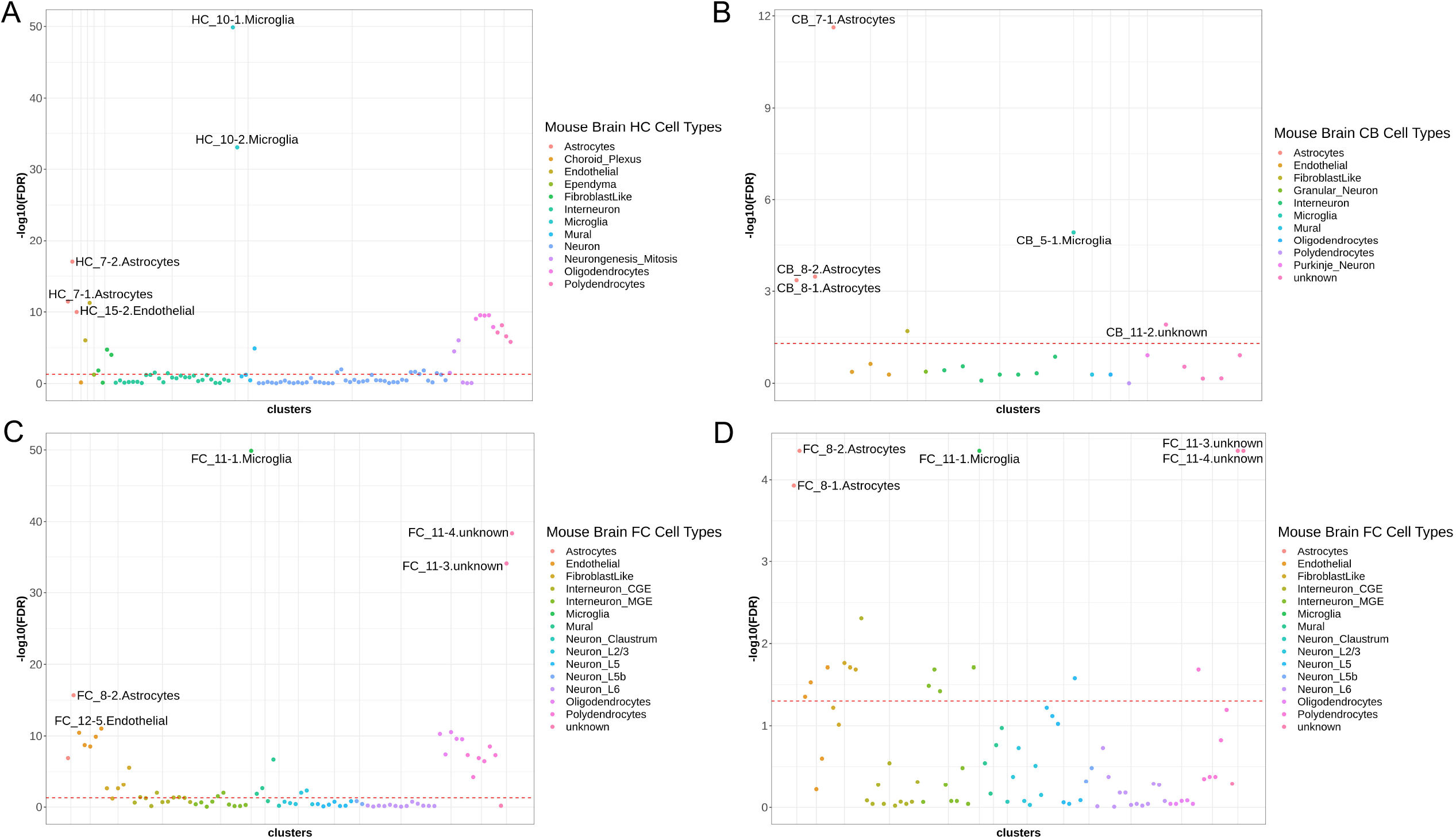
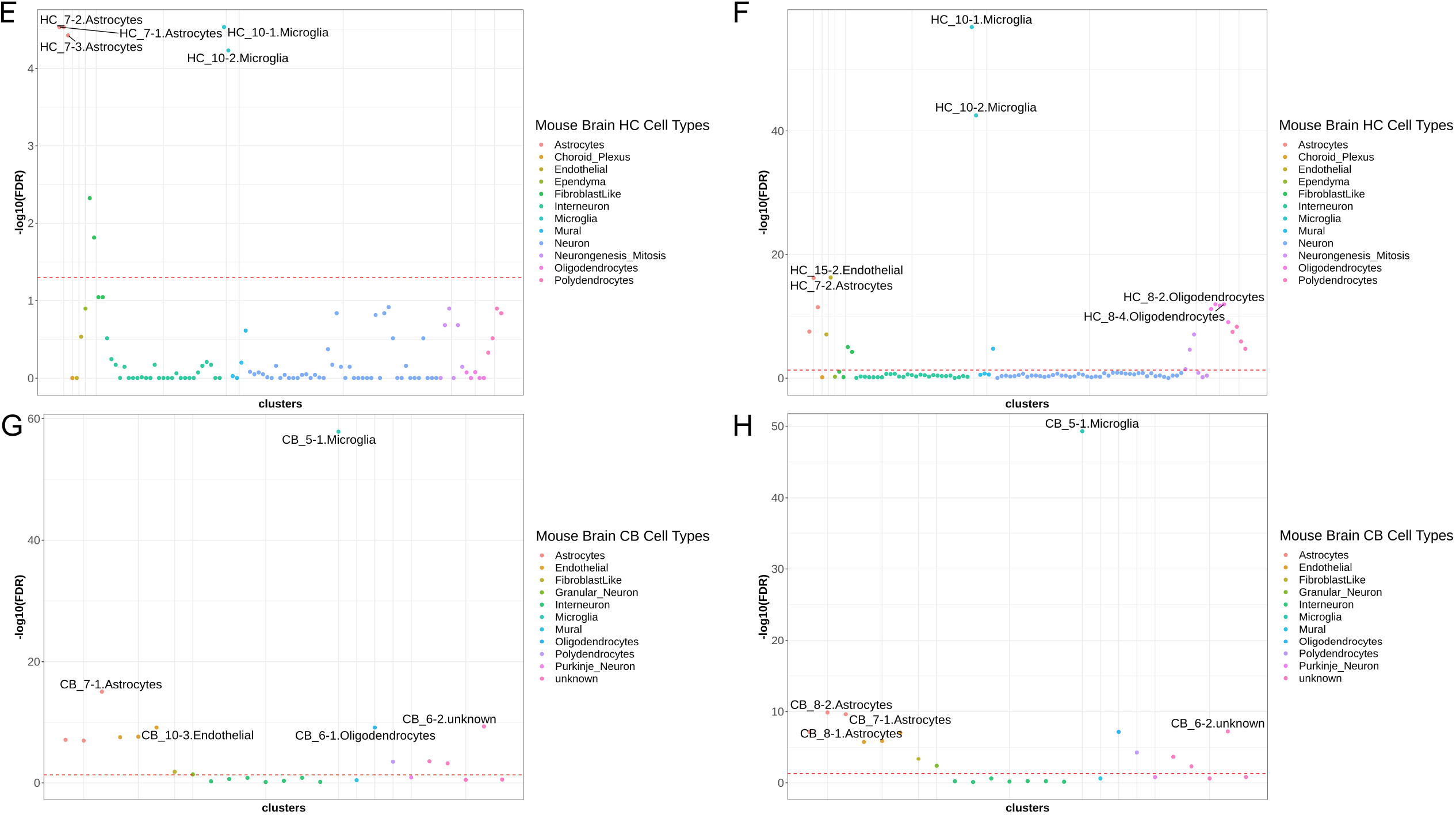
Comparison of *LRcell* analysis results when the tissue from which the DE gene lists is obtained match or un-match the tissue type from where the marker genes are collected. *LRcell* result when: (A) Both DE gene list and marker genes are obtained from HC. (B) Both DE gene list and marker genes are obtained from CB. (C) DE gene list obtained from HC and marker genes collected from FC. (D) DE gene list obtained from CB and marker genes collected from FC. (E) DE gene list obtained from CB and marker genes collected from HC. (F) DE gene list obtained from FC and marker genes collected from HC. (G) DE gene list obtained from FC and marker genes collected from CB. (H) DE gene list obtained from HC and marker genes collected from CB. Compared to (D) and (E), which analyze CB bulk DE genes, (B) has a higher resolution as the FDR ranges from 0 to 12.

